# Analysis of the Distal Urinary Tract in Larval and Adult Zebrafish Reveals Unrecognized Homology to the Human System

**DOI:** 10.1101/2023.01.30.526174

**Authors:** Ibrahim Jubber, Duncan R Morhardt, Jonathan Griffin, Marcus Cumberbatch, Maggie Glover, Yang Zheng, Ishtiaq Rehman, Catherine Loynes, Syed A Hussain, Stephen A Renshaw, Steven D Leach, Vincent T Cunliffe, James WF Catto

## Abstract

Little is known about the distal excretory component of the urinary tract in *Danio rerio* (zebrafish). This component is affected by many human diseases and disorders of development. Here, we undertook multi-level analyses to determine the structure and composition of the distal urinary tract in the zebrafish. *In silico* searches identified *uroplakin 1a, 2-like* and *3-like* genes in the zebrafish genome (orthologues to genes that encode proteins that characterise the urothelium in humans). *In situ* hybridization demonstrated *uroplakin-1a* expression in the zebrafish pronephros and cloaca from 96 hours post-fertilisation. H&E sectioning of adult zebrafish demonstrated two mesonephric ducts uniting into a urinary bladder which leads to a distinct urethral opening. Immunohistochemistry identified Uroplakin 1a and 2 expression within the urothelial luminal surface and Gata3 expression in cell layers that match human urothelial expression. Fluorescent dye injections demonstrated urinary bladder function, including urine storage, intermittent micturition, and a urethral orifice separate from the larger anal canal and rectum. Our findings reveal homology between the urinary tracts of zebrafish and humans and offer the former as a model system to study disease.

**Summary Statement:** The excretory components of the distal urinary tract in larval and adult zebrafish have been incompletely evaluated. Here, we demonstrate close homology in urinary tract anatomy between zebrafish and human, including the presence of a distinct adult urinary bladder capable of periodic micturition

## Introduction

The urinary tract is essential for the regulation of extracellular fluid and the excretion of metabolites. In humans, the urinary tract is composed of a multifunctional kidney, with excretory, metabolic, and endocrine roles, and an excretory component comprised of propulsive drainage tubes (ureters) leading to a contractile expansive storage vesicle (urinary bladder) with intermittent outflow via the urethra under voluntary and involuntary neurological control. The embryological development of the urinary tract is characterised by complex interactions between various embryological tissues.

In humans, the distal excretory component of the urinary tract begins development at approximately 4 weeks gestation. The cloaca is partitioned by the urorectal septum to form the ventral urogenital sinus and the dorsal anorectal canal. The urogenital sinus develops into the urinary bladder and urethra. The mesonephric ducts fuse with the urogenital sinus and the ureteric bud develops as an outgrowth from the distal mesonephric duct. The ureteric bud is involved in reciprocal induction events with the metanephric mesenchyme giving rise to the collecting ducts, renal pelvic and ureters. The distal mesonephric duct and ureteric bud undergo a remodelling process whereby the ureters fuse with the urogenital sinus and subsequently migrate superiorly and laterally to form the vesicoureteric junctions and the trigone. The distal mesonephric ducts migrate inferiorly to empty into prostatic urethra and forms the ejaculatory duct (Liaw et al., 2018).

The resultant excretory system is lined by urothelium, a multi-layered epithelium with a terminally differentiated superficial layer as well as intermediate and basal cell compartments (Dalghi et al., 2020). The superficial layer is composed of ‘umbrella cells’ which are characterised by the expression of uroplakin proteins. Uroplakin proteins are from a superfamily of tetraspanins and highly evolutionarily conserved across species (Garcia-Espana et al., 2006). They form an asymmetric unit membrane on the apical surface of the superficial urothelial layer, which is believed to have barrier function. Basal cell layers express transcription factor tumour protein 63 (TP63 or p63) (Steurer et al., 2021) and high molecular weight cytokeratins such as cytokeratin 5 (KRT5) (Volkel et al., 2022).

*Danio rerio* (zebrafish) models have been used to study the biology of the kidney. The zebrafish embryonic and larval kidney (pronephros) is comprised of two crescent shaped nephrons fused in the midline that course from the glomerulus to the pronephric ducts and cloaca (Naylor and Davidson, 2017). The mesonephros (the permanent adult kidney structure in zebrafish) develops at around 10 days post fertilisation (dpf) during the transition from larva to juvenile as the osmoregulatory demands of the fish increase (Diep et al., 2015). Whilst the zebrafish kidneys have been well characterised, the distal excretory component of the urinary tract is not well studied and it is unclear whether a bladder and urethra exist and exit the body separately from the intestinal tract (Holden et al., 2013; Outtandy et al., 2019).

Here we describe the zebrafish distal excretory urinary tract. We identify features similar to those observed in larger teleosts and mammals, including a urinary bladder that leads to a distinct urethra transporting urine externally. We also demonstrate intermittent micturition functionally *in vivo*.

## Results

### Uroplakin genes in the zebrafish

We searched for conserved uroplakin gene orthologues within the zebrafish genome. We identified three candidate genes: *uroplakin-1a* (ENSDARG00000021866.9), *uroplakin-2-like* (ENSDARG00000103164.2) and *uroplakin-3-like* (ENSDARG00000092872.2). Protein alignment revealed structural homology and inter-species conservation (Supplementary figures 1-3). Uroplakin-1a is a member of the tetraspanin superfamily of transmembrane proteins, which have 4 transmembrane helices (3 clustered towards the N terminus and 1 at the C terminus). Uroplakin-2 and 3 have a single transmembrane domain towards the C terminus. Mammalian Uroplakin-1a and 1b interact with Uroplakin-2 or 3a to form two heterodimers that can subsequently exit the endoplasmic reticulum. The two components of this dimer (1a/1b and 2/3a) function together as a unit (Garcia-Espana et al., 2006).

### *Uroplakin-1a* mRNA expression in zebrafish larval pronephros

Uroplakin-1a is the most highly conserved uroplakin and the only member present in zebrafish from the Uroplakin-1a/1b component of the dimer. Using *in situ* hybridisation, we identified the onset of *uroplakin 1a* expression to be around 96 hpf (hours post fertilization) in the zebrafish larval pronephros (Figure 1).

**Figure 1.**
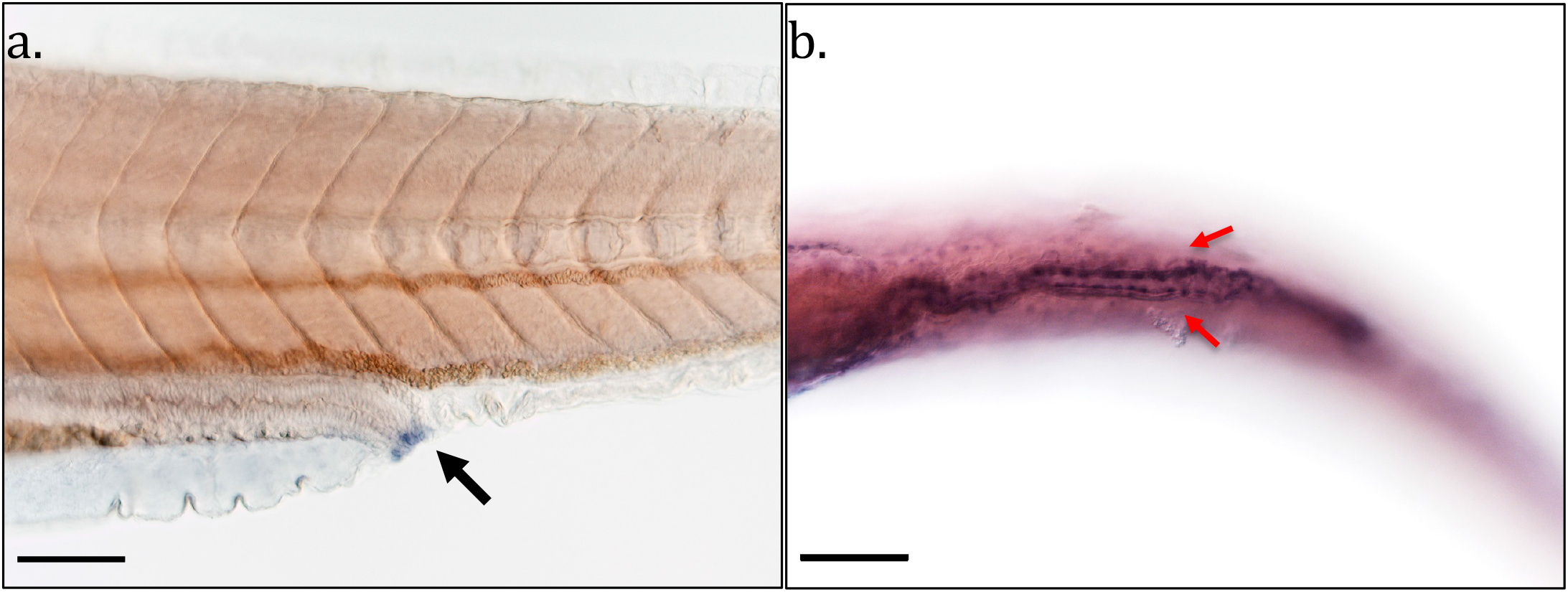
Expression of *uroplakin 1a* in Zebrafish Larvae Using *In Situ* Hybridisation. **A.** *uroplakin-1a* expression in the cloaca (Black arrow) at 96 hours post fertilisation (hpf) in the lateral view. **B.** *uroplakin-1a* expression in the two pronephric ducts (red arrows) as they drain from the kidney towards the cloaca at 120 hpf in the ventral view. Scale bars=50μm.

### Adult zebrafish demonstrate conserved anatomy of the excretory urinary tract

To explore anatomical structure, we sectioned and stained adult zebrafish from the kidney to the cloacal region in transverse, coronal and sagittal orientations. Microscopy revealed two mesonephric ducts leaving the distal kidney, traveling inferiorly and caudally before fusing into a single collecting vesicle, with features of a urinary bladder (Figure 2). In contrast to the human urinary bladder urothelium which is multiple cell layers in thickness, the zebrafish urinary bladder epithelium is mostly one to two cells thick. The innermost cells of the zebrafish urinary bladder epithelium (in contact with the lumen) have a similar appearance to the outermost cells.

**Figure 2.**
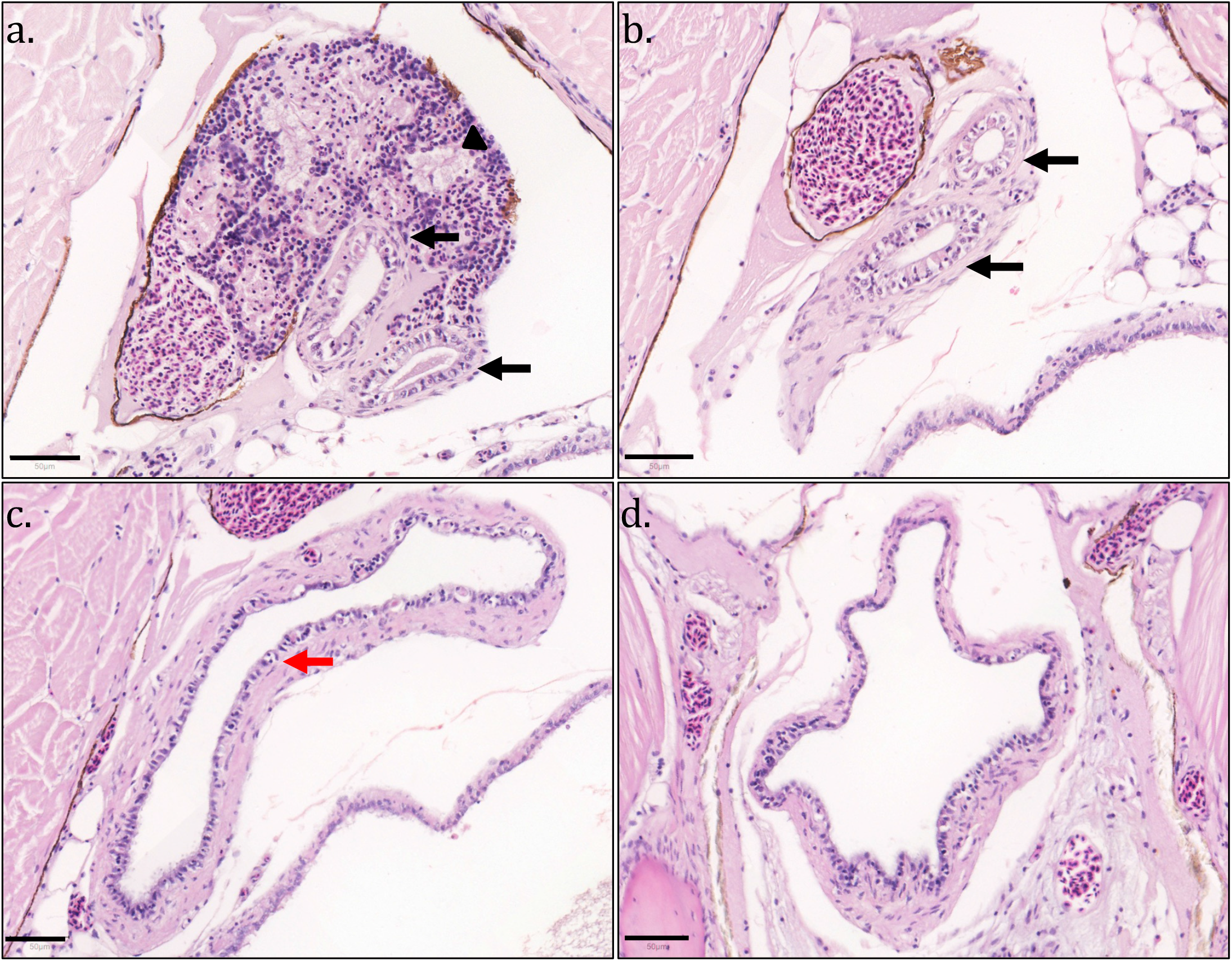
Anatomy of Adult Zebrafish Mesonephric Ducts and Urinary Bladder. H&E- stained adult zebrafish sections in transverse orientation from cranial to caudal. **A.** Kidney (black triangle) converges onto two mesonephric ducts (Black arrows). **B.** Two mesonephric ducts (black arrows) distal to kidney in retroperitoneum. **C.** The two mesonephric ducts unite into a single urinary bladder lined by epithelial cells (red arrow). **D.** Inferior aspect of urinary bladder. Scale bars=50μm.

The urinary bladder leads to a single tube that is separate from the anal canal and rectum in keeping with a urethra. In females, the urethra and oviduct are separate tubes throughout their lengths. However, in males the ejaculatory duct joins into the urethra, enabling ejaculation to occur via the urethra. The urethra in both males and females is lined by an epithelial layer which is surrounded by a deeper connective tissue layer rich in collagen (Figure 3).

**Figure 3.**
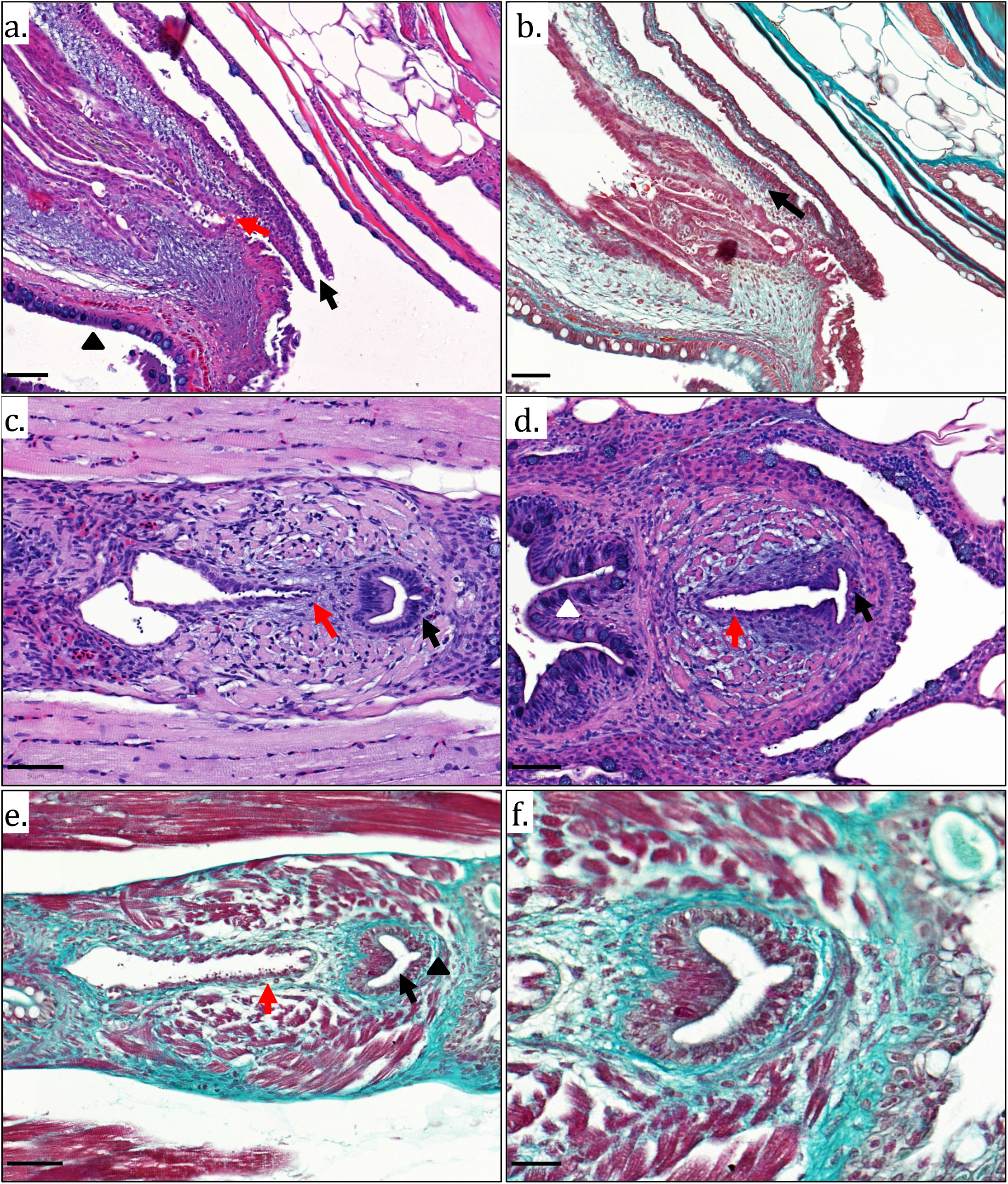
Anatomy of Adult Zebrafish Urethra. A. H&E stain (sagittal orientation) of adult female zebrafish cloacal region demonstrating distinct and separate urethral orifice (black arrow), oviduct (red arrow) and rectum (black triangle). B. Masson’s trichrome demonstrating connective tissue (light green colouration, black arrow) surrounding female urethra. C. H&E stain (coronal orientation) of adult male zebrafish cloacal region demonstrating urethra (black arrow), ejaculatory duct (red arrow). D. H&E stain (coronal orientation) of adult male zebrafish cloacal region demonstrating ejaculatory duct (red arrow) opening into urethra (black arrow) and distinct separate rectum (white arrow). E. Masson’s trichrome stain of male adult cloacal region (coronal orientation) demonstrating muscle fibres surrounding ejaculatory duct (red arrow) and connective tissue (light green colouration, black triangle) surrounding urethra (black arrow). F. Masson’s trichrome stain of male zebrafish urethra at higher magnification Scale bars =50μm for A-E and 20μm for F.

To understand protein conservation, we determined the expression of various proteins that characterise the human urothelium. GATA3 is a pioneer transcription factor and urothelial differentiation marker that is highly expressed in normal human urothelium (Miettinen et al., 2014). Immunohistochemistry revealed strong nuclear GATA3 expression in the zebrafish urothelium that matches human urothelial expression (Figure 4).

**Figure 4.**
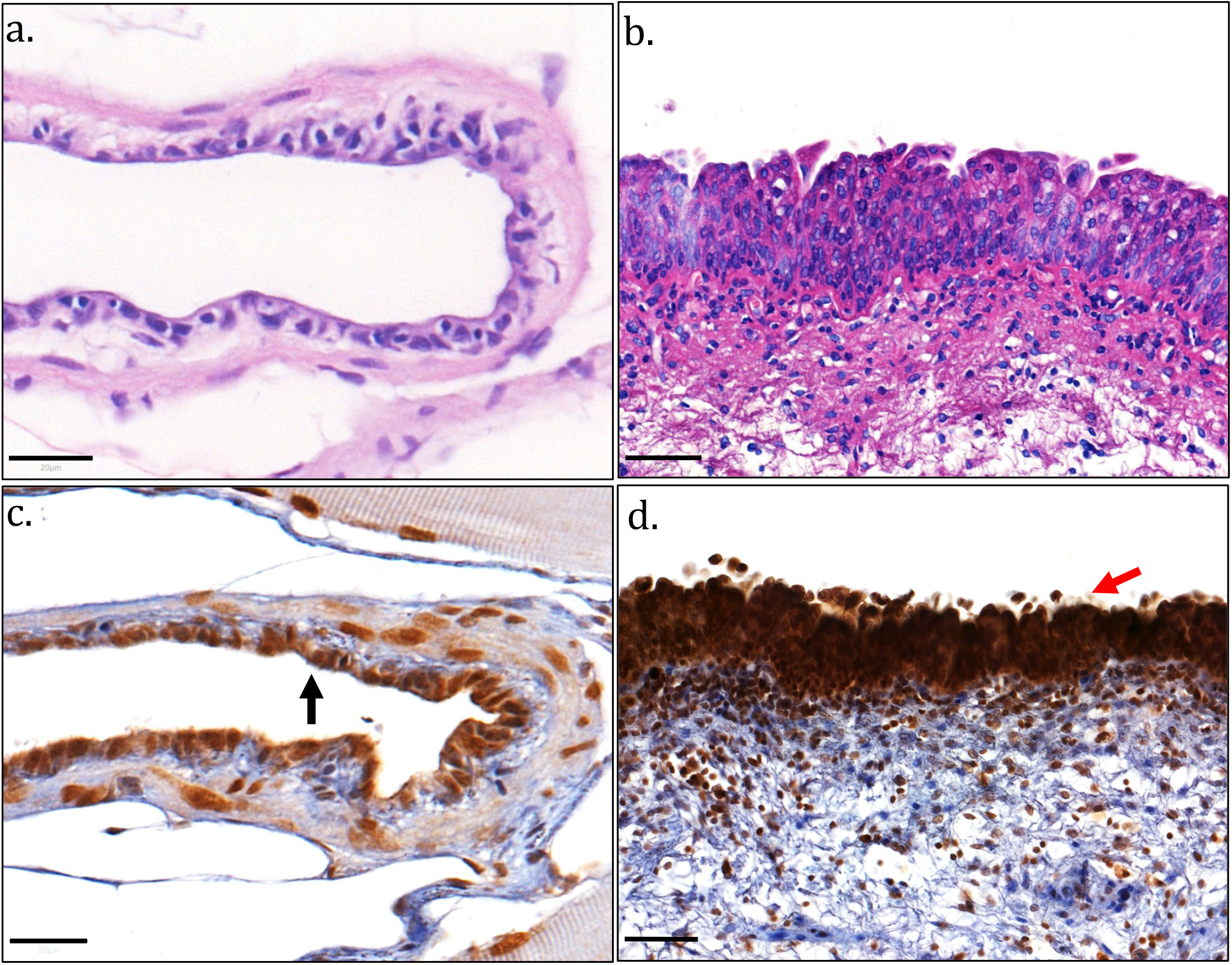
Expression of GATA3 in Zebrafish and Human Urinary Bladder Tissue Sections Using Immunohistochemistry. A. Hematoxylin and eosin stain of coronal zebrafish urinary bladder section and B. Human urinary bladder sections for reference. C. Nuclear expression of Gata3 in zebrafish urinary bladder urothelium (black arrow). D. Nuclear expression of GATA3 in human urinary bladder urothelium (red arrow). Scale bar=20μm for A and C. Scale bar=50μm for B and D.

Uroplakins are tetraspanin dimers that characterise the urothelium (Dalghi et al., 2020). Immunohistochemistry revealed cytoplasmic and membranous expression of Uroplakin-1a and Uroplakin-2l within the epithelium of the zebrafish urinary bladder and mesonephric duct, in a pattern similar to that seen in humans urothelium (Figure 5). Having demonstrated expression of proteins found in the superficial urothelial layer, we performed immunohistochemistry for known markers of human basal urothelial layers KRT5 (Volkel et al., 2022) and basal and stem cell marker CD44 (McKenney et al., 2001). Cytoplasmic KRT5 expression was seen in both the zebrafish urinary bladder and mesonephric duct epithelium, in similar patterns to the human tissues. The basal and stem-like marker CD44 showed cytoplasmic expression in the zebrafish urinary bladder urothelium in a similar manner to the human urinary bladder urothelium. There was a difference between the expression of KRT5 between zebrafish and human urothelium: they were both expressed throughout the zebrafish urothelium but only in the basal cell layer of the human urothelium (Figure 6).

**Figure 5.**
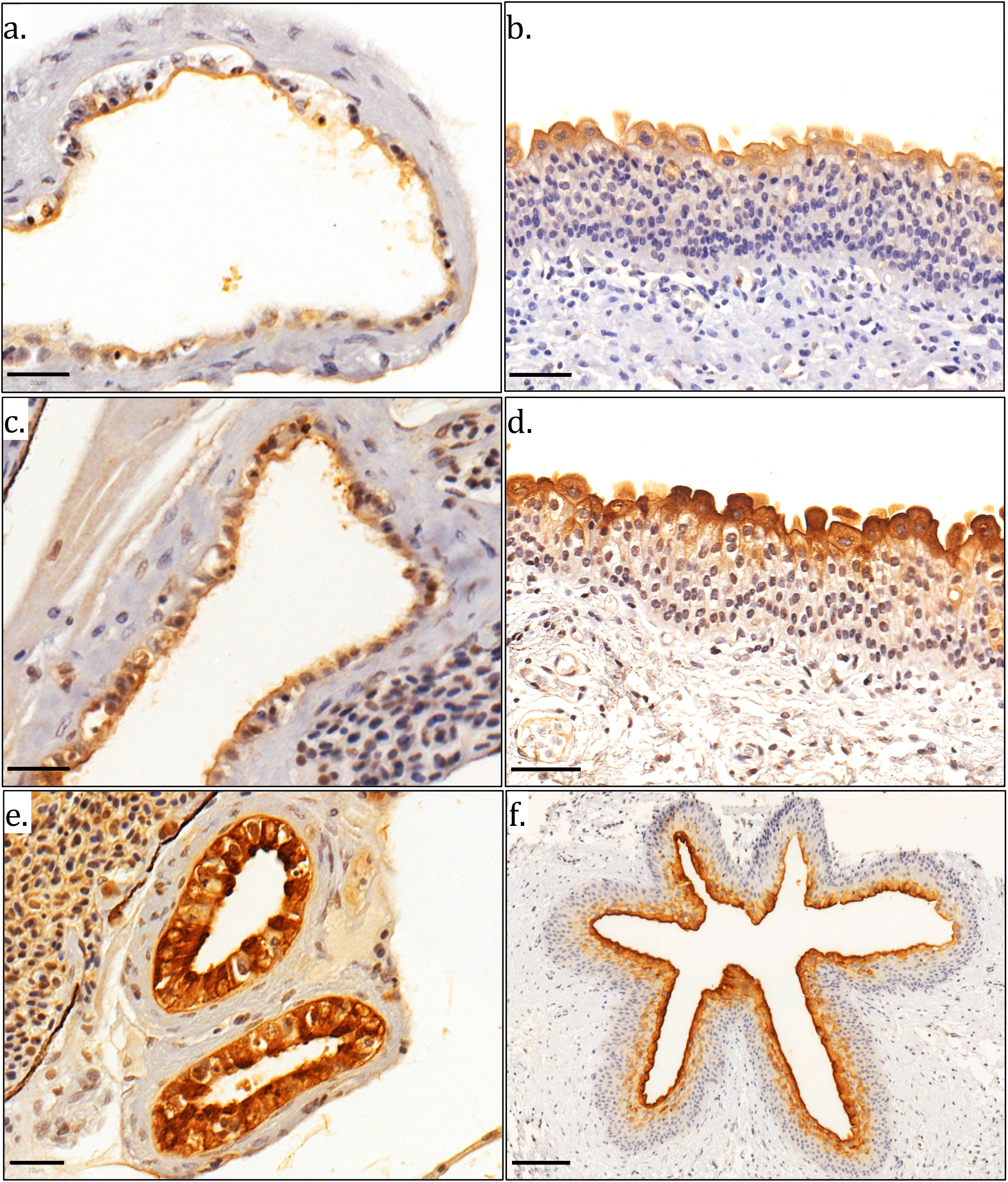
Expression of Uroplakins in Zebrafish and Human Urinary Bladder Tissue Sections Using Immunohistochemistry. A. Cytoplasmic and membranous expression of urothelium-specific protein Uroplakin 1a (Upk1a) in zebrafish urinary bladder urothelium. B. Cytoplasmic UPK1A expression in human urinary bladder urothelium. C. Uroplakin-2l (Upk2l) expression in zebrafish urinary bladder. D. UPK2 expression in human urinary bladder. E. Upk2l expression in zebrafish mesonephric duct epithelium. F. UPK2 expression in human ureter urothelium. Scale bar= 20μm for A, C & E. Scale bar=50μm for B & D. Scale bar= 100μm for F.

**Figure 6.**
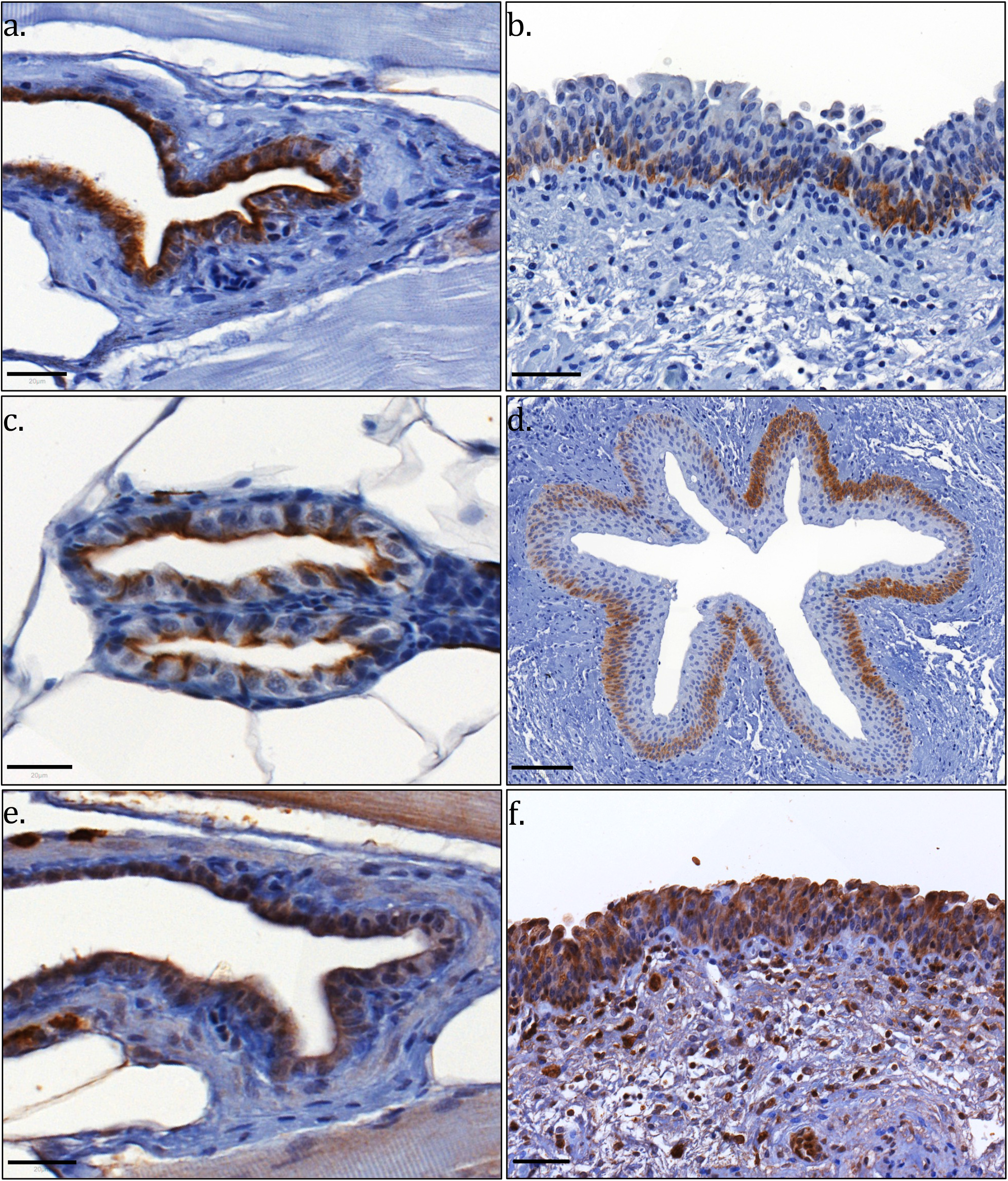
Expression of Urothelial Basal Cell Layer Markers in Zebrafish and Human Urinary Bladder Tissue Sections Using Immunohistochemistry. A. Cytoplasmic expression of Krt5 in zebrafish urinary bladder urothelium. B. Cytoplasmic KRT5 in human urinary bladder urothelium. C. Cytoplasmic Krt5 expression in zebrafish mesonephric duct epithelium. D. Cytoplasmic KRT5 expression in human ureter urothelium. E. Cytoplasmic CD44 expression in zebrafish urinary bladder urothelium. F. Cytoplasmic CD44 expression in human urinary bladder urothelium. Scale bar = 20μm for A, C, E. Scale bars=50μm for B, F. Scale bar=100μm for D.

Masson’s trichrome straining demonstrated that the wall of the zebrafish mesonephric ducts and urinary bladder have a connective tissue layer rich in collagen deep to the epithelial layer. Similar to the zebrafish mesonephric duct, the human ureter also as a connective tissue layer deep to the epithelial layer (Figure 7). However, the human ureter has a muscle layer deep to the connective tissue (lamina propria) layer which is not present in the zebrafish mesonephric duct. Masson’s trichrome staining demonstrated the zebrafish urinary bladder wall has, deep to the epithelial layer, a connective tissue layer and a deeper thin layer of muscle fibres. The structure of the human urinary bladder wall follows a similar pattern (Figure 7).

**Figure 7.**
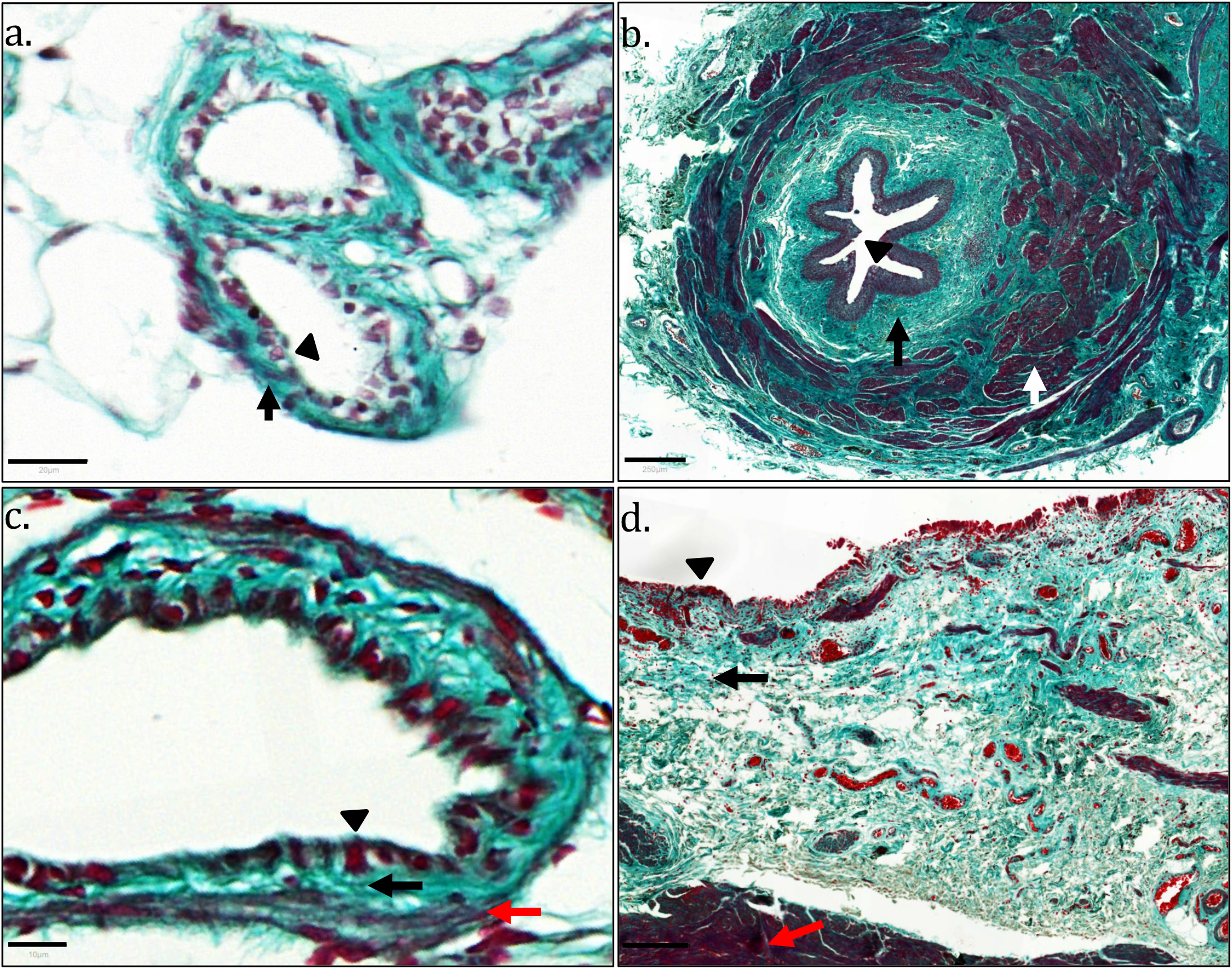
Structure of Zebrafish Mesonephric Duct and Urinary Bladder Wall. A. Masson’s trichrome stain of zebrafish mesonephric duct sections (coronal orientation) demonstrating connective tissue layer (light green colouration, black arrow) deep to epithelial layer (black triangle). B. Masson’s trichrome of human ureter section, for comparison, demonstrating connective tissue layer (light green colouration, black arrow) deep to epithelial layer (black triangle) and muscle layer (white arrow). C. Masson’s trichrome stain of zebrafish urinary bladder section (Coronal orientation) demonstrating, deep to epithelial layer (black triangle), a connective tissue layer (light green colouration, black arrow) and thin muscle fibres (red fibres, red arrow). D. Masson’s trichrome stain of human urinary bladder section demonstrating, deep to epithelial layer (black triangle), a connective tissue layer (light green colouration, black arrow) and a muscle layer (red fibres, red arrow). Scale bar=20 μm for A, 250 μm for B, 10 μm for C and 200 μm for D.

### In Vivo Analysis of Storage and Voiding Function in Adult zebrafish

To trace the course of urine in the excretory component of the zebrafish urinary tract, antegrade filling of the urinary system via pericardial injection and retrograde filling of the anal canal and rectum was performed with dextran-conjugated Alexa dyes on anaesthetised adult zebrafish (Figure 8). Cannulation of the anal canal and rectum for the retrograde studies was performed carefully using a 1μl syringe and approximately 0.2μl of dextran conjugated Alexa 568 was injected. The pericardial space was injected with Alexa 488. The fish were imaged on a confocal microscope over 10-20 minutes. By approximately 13 minutes after pericardial injection, urine (marked by green fluorescence) had accumulated in the urinary bladder region and micturition was demonstrated ventral to the urinary bladder (Figure 8a-c). Intermittent micturition continued for several minutes and demonstrated periods of pausing (Supplementary Video 1). Excreted urine was regularly washed away using a pipette in order to continually visualize the micturition process. Confocal images of the region of urinary expulsion (marked by pericardially injected Alexa dye) demonstrated the urethral orifice as a tubular structure separate from the anal canal opening (Figure 8d-f).

**Figure 8.**
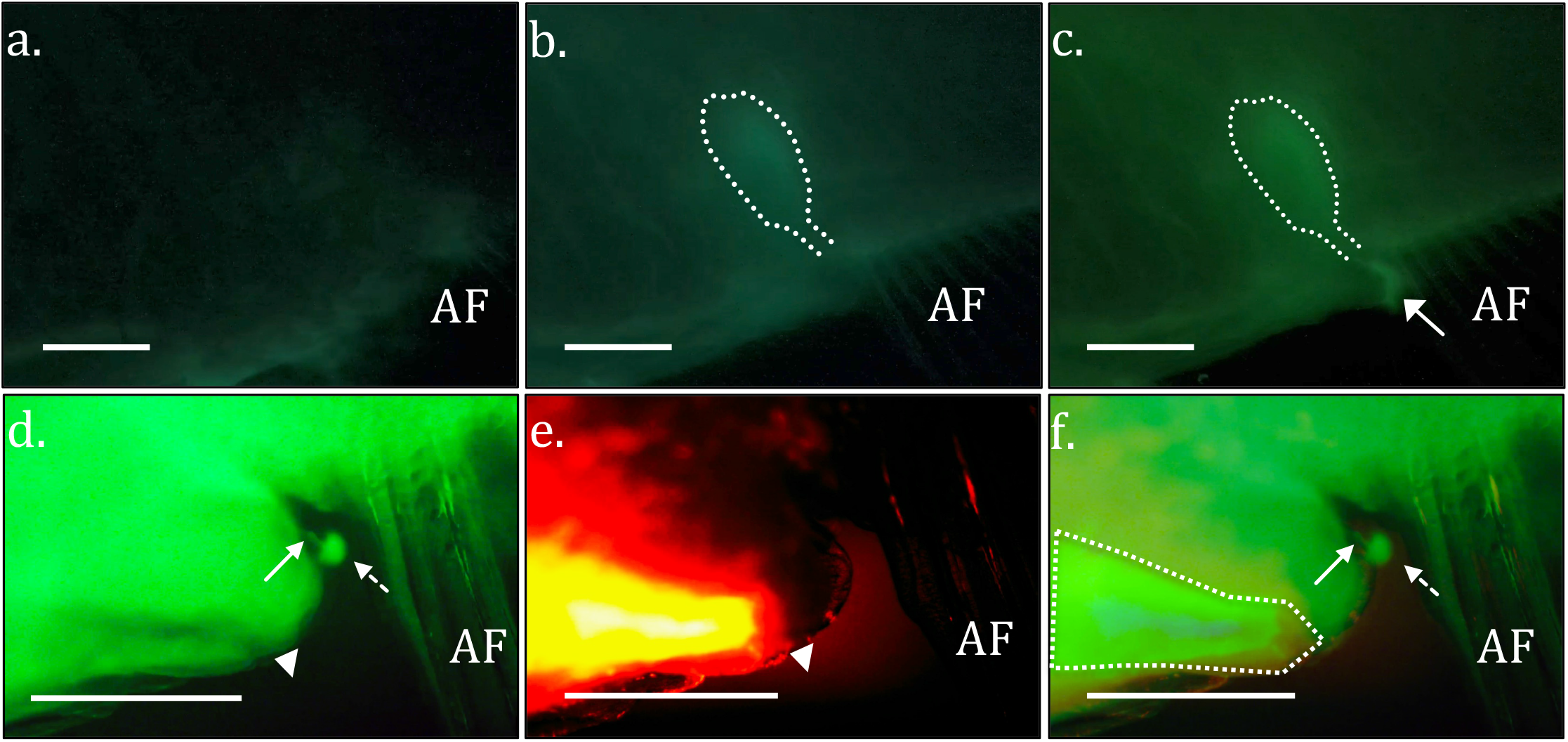
Antegrade and Retrograde Functional Urinary Studies in Live Adult Zebrafish. Alexa 488 conjugated dextran was injected into the pericardial space and pelvic region was imaged at A. <30 seconds, B. 10 minutes and C. 13 minutes post injection. Urinary bladder structure is outlined in white, and the expulsion of urine is marked with a white arrow. D. Epifluorescent imaging demonstrates urinary flow to a urethra (white arrow) posterior to the anal canal and rectum (white triangle). This urinary structure is unique from the anal canal and rectum and separately expels urine (white dotted arrow). E. Retrograde counterinjection of dextran-conjugated Alexa 568 into anal canal (white triangle) reveals outline of rectum in orange. F. Merged image identifies a urinary channel (white arrow) separate from the anal canal and rectum (outlined in white). Anal fin (AF) is identified for reference. Scale bar for A-C= 0.5 cm. Scale bar for D-H= 0.5 mm.

While the injection studies demonstrated a bladder structure and separate orifice, urine accumulation in a non-partitioned imaging dish prevented satisfactory quantification of urinary function. We therefore designed an independent, two-chambered apparatus allowing for continuous exposure to tricaine anaesthesia in the gill region, combined with continuous flushing of the excreted urine at a stable rate, thus permitting quantification of voiding function. Movies were analysed to first identify the correlation of bladder emptying with urinary excretion. Fluorescent quantification of dye demonstrates that episodes of bladder emptying coincide with high volume voiding via the urethral meatus (Figure 9a-b). Average bladder emptying interval (E.I.) was 302 ± 37 (S.E.M) seconds (n=4 animals, 13 E.Is, Figure 9c). The mechanism of emptying was further investigated by examining sequential segments of the bladder structure. Minor contractions were seen in the bladder during filling, but bladder emptying regularly occurred with a concerted contraction which involved simultaneous emptying of each segment of bladder (proximal, middle, and distal, Figure 9d). Peristaltic emptying, where sequential segments propel urine along the bladder to excrete, was not appreciated. In addition to well-correlated large voids, smaller or more frequent voids were also occasionally observed without significant emptying.

**Figure 9:**
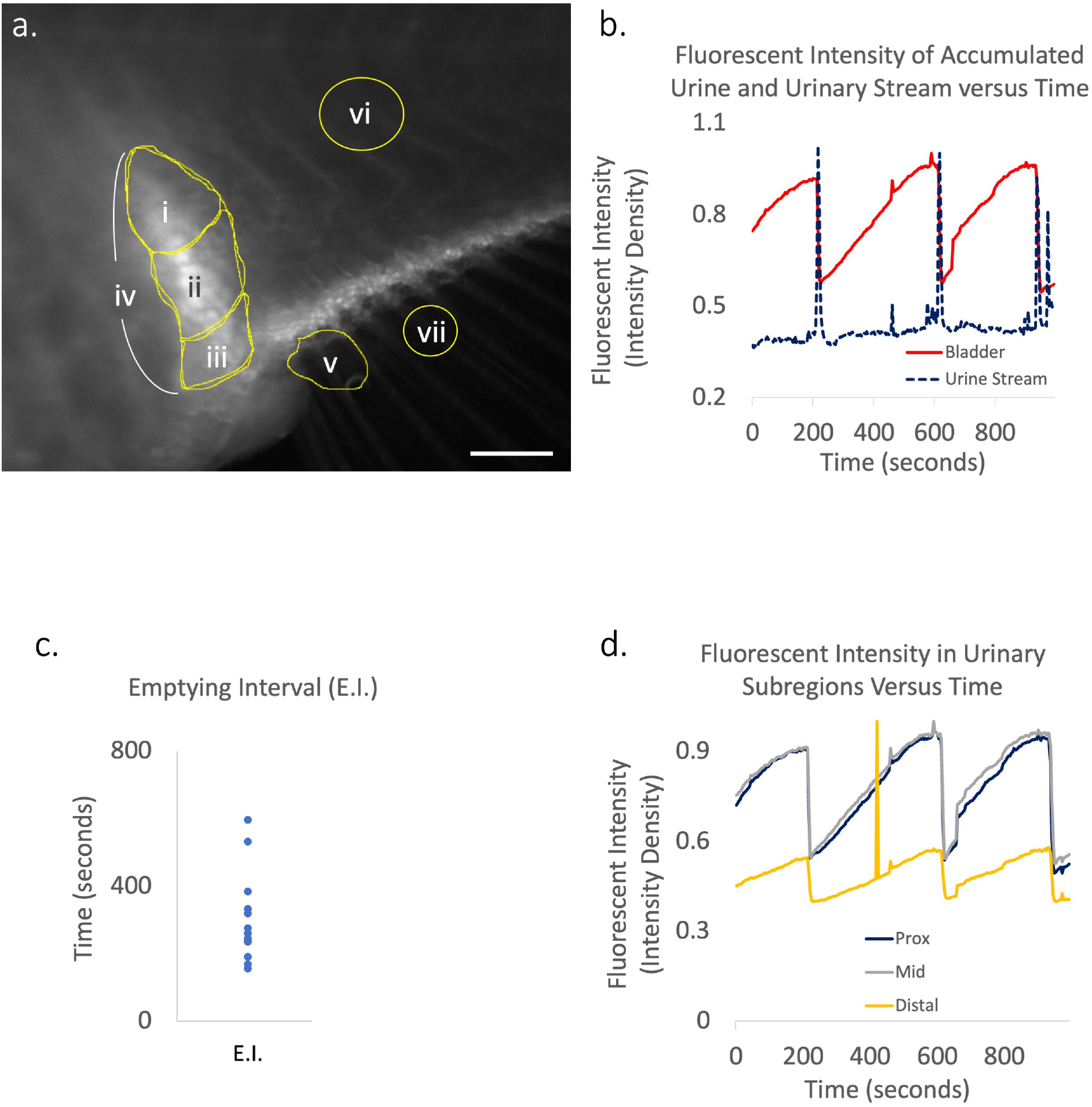
Quantification of Urinary Excretion in Zebrafish. A. After pericardial Alexa 488 injections, serial images were taken of zebrafish urinary structures (scale bar = 0.4mm). Regions of interest were identified as i, proximal bladder, ii, middle bladder, iii, distal bladder, iv, total bladder, and v, urinary stream. vi and vii were selected for normalization of body and urinary streams, respectively. B. Plots of total bladder(iv) fluorescence intensity density were plotted alongside urinary stream (v) intensity over time. C. Distribution of bladder emptying interval (E.I) in seconds were plotted. D. Bladder subregion fluorescent intensity is plotted against time revealing simultaneous contraction, and not sequential peristalsis, empties the full urinary bladder.

## Discussion

This study characterised the distal excretory component of the urinary tract in both larval and adult zebrafish and demonstrated close homology in distal urinary tract anatomy and function between zebrafish and human. We demonstrate the presence of uroplakin 1a, a characteristic urothelial protein, in the zebrafish embryo pronephros starting at 96 hpf. Secondly, we characterised key components of the adult zebrafish urinary excretory system and identified a urinary bladder structure, arising from the fusion of the two mesonephric ducts. Thirdly, this study demonstrated the zebrafish urinary bladder empties via a distinct urethra. Lastly, we demonstrated functionally in vivo in anaesthetised adult zebrafish that urine accumulates in the bladder and is intermittently released.

Using whole mount *in situ* hybridisation, we detected the presence of *uroplakin1a* expression in the zebrafish larval pronephric tubules and ducts at 96 hpf and 120 hpf. This is supported by the results of a previous study demonstrating expression of another uroplakin protein, Upk3l, at the apical surface of the pronephric tubules and ducts (Mitra et al., 2012). Morpholinos against *upk3l* led to embryos with reduced pronephric function believed to be due to defects in epithelial polarity and morphogenesis. The zebrafish pronephric epithelium and the human urothelium both play roles in urine excretion function and are in direct contact with urine. Our findings are suggestive of structural homology between these two structures. Uroplakins are transmembrane proteins that form an as an asymmetric unit membrane (AUM) on the apical surface of the superficial layer of human urothelium (Dalghi et al., 2020). The function of uroplakins is not entirely clear. Mouse knockout studies suggest that they play a role in apical membrane permeability barrier function of urothelium (Hu et al., 2000; Hu et al., 2002). Abnormal uroplakins have also been linked to urinary tract malformations in mouse knockout and human genetic studies (Jenkins et al., 2005; Kong et al., 2004).

We demonstrated that zebrafish possess two mesonephric ducts that arise from the kidneys and unite into a urinary bladder. The zebrafish mesonephric ducts and urinary bladder are lined by an epithelium that expresses proteins characteristic of human urothelium suggesting structural similarity and evolutionary conservation. These are important findings as previously there have been conflicting reports regarding the presence of a urinary bladder in zebrafish (Baranowska Körberg et al., 2015; Holden et al., 2013; Outtandy et al., 2019). However, the presence of mesonephric ducts and urinary bladder has been described in other teleost species (Curtis and Wood, 1991; Macrì et al., 2014). We did note two main differences between the zebrafish urinary epithelium and human urothelium. Firstly, in comparison to the morphology of the human urinary bladder urothelium which has multiple cell layers, the zebrafish urinary bladder epithelium is one to two cell layers thick. Secondly, human urothelium is composed of superficial (‘umbrella’), intermediate, and basal cell layers which are characterised by expression of different proteins in the different layers (Dalghi et al., 2020). Although the zebrafish urinary bladder epithelium is one two cells thick, we did demonstrate that the zebrafish urinary bladder epithelium expresses both superficial markers (Uroplakins) and basal markers (Cytokeratin 5 and Cd44).

This study also showed structural similarity in the lower urinary tracts distal to the urinary bladder in zebrafish and humans. We demonstrated that the zebrafish urinary bladder leads to a urethra and urethral orifice that is separate from the hindgut and surrounded by a deeper connective tissue layer rich in collagen. These structures have not previously been described in zebrafish. Furthermore, we demonstrated sex differences in urethral anatomy in zebrafish by demonstrating that the male zebrafish ejaculatory duct joins the urethra (similar to humans) whereas the female oviduct is a separate structure to the urethra.

In anaesthetised adult zebrafish, we showed in vivo that urine is stored and intermittently released. Intermittent release of urine has been reported in another larger teleost species, the freshwater rainbow trout where urine is stored for 25-30 minutes (Curtis and Wood, 1991). Urine storage in the freshwater rainbow trout enables solute reabsorption (sodium and chloride ions) to take place functioning as an adjunct to kidney function (Curtis and Wood, 1991). Although, this is not considered to be a traditional function of the mammalian urinary bladder, recent data in a porcine model is supportive of urinary bladder reabsorption function in mammals (Manso et al., 2019). Urine storage in the freshwater rainbow trout has also been suggested as potentially important for survival by avoiding the persistent release of olfactory signals to potential nearby predators (Curtis and Wood, 1991).

## Conclusion

The adult zebrafish distal urinary excretory system shares several features with its human counterpart. The zebrafish system has two mesonephric ducts leading to a urinary bladder lined by a uroplakin expressing urothelium. The urinary bladder, in turn, leads to a urethra and urethral orifice. These findings present an opportunity to model human lower urinary tract diseases in zebrafish.

## Materials and Methods

### Genome alignment

Genomic DNA and protein sequences for uroplakin genes were identified in humans (Homo sapiens), mouse (Mus musculus), cow (Bos taurus) and the zebrafish by searching the NBCI genome and protein browser. Protein sequences were aligned using the muscle tool online using Snapgene software (Version 4.3.11).

### Zebrafish Maintenance and Strains

Zebrafish were raised and housed in the Bateson Centre at the University of Sheffield in aquaria approved by the UK Home Office and maintained according to standard protocols (maintained at 28.5 degrees) in accordance with the UK Animals (Scientific Procedures) Act 1986. Wild type zebrafish were from stocks held at the Sheffield Biological Services Aquaria.

### Histology

To obtain zebrafish paraffin tissue sections, adult fish were culled and subsequently fixed 10% Formalin for 3-4 days at 4 degrees or 2 days at room temperature. The samples were decalcified using 0.25M EDTA pH 8 at room temperature for 2-4 days and then immersed in 70% ethanol. The samples were dehydrated through ascending concentrations of ethanol (70% to 100%) and transferred to xylene, before infiltration and embedding in paraffin wax. A microtome was used to section wax embedded fish at 3.5-5 μm sections. Fish samples were either oriented transversely, coronally or sagittally. Human bladder tissue was obtained with ethical approval the South Yorkshire Research Ethics Committee (REC reference number: 10/H1310/73). For haematoxylin and eosin (H&E) staining, tissue sections were dewaxed with xylene and rehydrated in descending concentrations of ethanol to water before staining.

For Masson’s trichrome staining, sections were dewaxed in xylene and rehydrated in descending concentrations of ethanol to water and subsequently mordanted in Bouins solution for 1 hour at 56 degrees. The sections were then stained with Wiegert’s haematoxylin, Ponceau fuschin and light green SF. Differentiation was performed using phosphomolybdic acid solution prior to light green stain.

### Immunohistochemistry

Sections were dewaxed and rehydrated. Endogenous peroxidase was blocked with 3% hydrogen peroxide/methanol solution (one part 30% hydrogen peroxide and nine parts methanol) at room temperature. The Avidin and biotin blocking kit (Vector Laboratories now called 2b Scientific) was used (for Krt5, Gata3 and Cd44 markers) to block signal from endogenous biotin, biotin receptors and avidin binding sites present in tissues. 10% serum block was used to reduce non-specific binding (rabbit and goat sera). The following antigen retrieval methods were used: 0.01 M Citrate Buffer (heated in microwave) and DAKO Target Retrieval Solution, S1699 (heated in antigen retriever).

The following primary antibodies were used: Uroplakin 1a (N-16):sc-15170 (Goat polyclonal) at 1:400 dilution for zebrafish tissue and 1:200 dilution for human tissue (Santa Cruz Biotechnology), Uroplakin 2 (k-18): sc-15179 (Goat polyclonal) at 1:400 dilution for zebrafish tissue and 1:100-1:400 dilution for human tissue (Santa Cruz Biotechnology), Cytokeratin 5/6 (mouse monoclonal) at 1:100 to 1:200 dilution for zebrafish tissue and 1:100 dilution for human tissue (Dako), GATA 3 (SAB2100898) (rabbit) 1:400 dilution for zebrafish tissue and 1:100 dilution for human tissue (Sigma Aldrich) and and CD44 (SAB1405590) (mouse) at 1:400 dilution for zebrafish tissue and 1:50 dilution for human tissue (Sigma Aldrich). The following secondary antibodies were used: Biotinylated Rabbit anti-goat 1:200 dilution (Vector Laboratories now 2b Scientific), Biotinylated Goat anti-rabbit 1:200 dilution (Vector Laboratories now 2b Scientific), Biotinylated Goat anti-mouse 1:200 dilution (Vector Laboratories now 2b Scientific). Vectastain Elite ABC kit, Peroxidase (Vector Laboratories now 2b Scientific) and DAB (3,3’-diaminobenzidine) substrate kit, Peroxidase (Vector Laboratories now 2b Scientific) were used for detection.

### Whole mount In situ hybridization

For Digoxygenin-labelled RNA probe production, the probe for *uroplakin 1a* was synthesised from linearised plasmid DNA obtained from a plasmid vector containing the coding sequence. The coding sequence was contained in the: p-express 1 plasmid. Details of plasmids used to synthesise RNA probes are available on request. For the anti-sense *uroplakin 1a* probe, the p-express 1 plasmid was linearised using the EcoRI restriction enzyme and the T7 RNA polymerase enzyme was used for transcription. For the sense *uroplakin 1a* probe, the p-express 1 plasmid was linearised using the NOTI restriction enzyme and the SP6 RNA polymerase enzyme was used for transcription. Whole-mount *in situ* hybridisation was performed using standard procedures (Westerfield, 2000).

### Anatomical Tracing Studies

Adult fish were anesthetized with tricaine solution. Dextran-conjugated Alexa dyes (Thermo-Fisher, Waltham, MA) were prepared in phosphate buffered saline and had a concentration of 10-20ug/ul. Using 1μl syringes, the anal canal and rectum was carefully cannulated and injected ~0.2ul Alexa 568 and the pericardial space injected with Alexa 488. Fish were imaged over 10-20 minutes and maintained under anaesthesia with flushes of tricaine solution. For urinary function studies, fish were similarly anesthetized in titrated tricaine and only intracardiac injection was performed. Fish were placed in a partitioned custom physiology chamber to allow for differential anaesthetic and flushing flow. Images were obtained every four seconds. Fish were sacrificed in ice water baths after imaging.

### Imaging and Software

A Leica M165 FC (Leica Microsystems) microscope was used for screening fluorescent microscopy. The ISH images were obtained using Leica Microsystems DFC 420 C1.8 microscope. Anatomical tracing studies was performed with a Zeiss Axiocam 208 on a fluorescent Leica stereoscope. Images were captured with Labscope software (Zeiss, Germany). In vivo urinary studies were performed using the Zeiss AxioZoom steromicroscope and Zen software. Images were then selected for regions of interest, as identified in Figure 9a, and serially measured for intensity density in FIJI which were plotted against time. Confocal images were obtained on an Andor Spinning disk confocal microscope. Movies clips were edited using iMovie software (Apple, Cupertino, CA). The Panoramic 250 Flash III slide scanner (3DHISTECH) was used to image the histology slides. Histology images were processed using Qupath version 0.1.2 (Bankhead et al., 2017).

## Acknowledgements

We thank Dr Nick Van Hateran, Dr Darren Robinson and the Wolfson Light Microscopy Facility at the University of Sheffield, UK. We also thank Life Sciences Light Microscopy Facility, Dartmouth College, USA.

## Figure Legends

**Supplementary Figure 1. Alignment of Human, Mouse and Zebrafish Uroplakin 1a Amino Acid Sequences.** Muscle Tool used for alignment. Human sequence used as reference. Alignment highlighted in green.

Supplementary Figure 2. Alignment of Human, Mouse and Zebrafish Uroplakin 2 Amino Acid Sequences. Muscle Tool used for alignment. Human sequence used as reference. Alignment highlighted in green.

**Supplementary Figure 3. Alignment of Human, Mouse and Zebrafish Uroplakin 3a Amino Acid Sequences.** Muscle Tool used for alignment. Human sequence used as reference. Alignment highlighted in green.

**Supplementary Figure 4. Sense Control for *Uroplakin 1a In Situ* Hybridisation in Zebrafish larvae at 96 hpf.** Scale bar= 50μm.

**Supplementary Figure 5. Negative (No Primary Antibody) Controls for Immunohistochemistry.** A. No primary antibody control for biotinylated goat anti-rabbit secondary antibody, B. No primary antibody control for biotinylated rabbit anti-goat secondary antibody. C. No primary antibody control for biotinylated goat anti-mouse secondary antibody. Scale bars= 50μm.

**Supplementary Video 1: Micturition in Zebrafish.** Pericardial injections were performed in adult fish as described. Counter injection into the cloacal channel was performed with dextran-conjugated Alexa 568. Separate channels were obtained by alternating prisms. Within 10 minutes, urine was expelled from the dorsal urethral meatus. Excreted urine was regularly washed away using a pipette in order to continually visualize the micturition process. Timescale is approximately 16x acquired speed. Images have same size scale as images in Figure 8d-f.

**Supplementary Video 2: Bladder emptying with voiding activity.** Pericardial injections were performed as described. Gentle flow of tank water removed excreted dye that was accumulated. Serial images were taken every 4 seconds. Movie is run at 32x speed. Scale bar=1mm.

