## Supplementary Figures for "Analysis of the Distal Urinary Tract in Larval and Adult Zebrafish Reveals Unrecognized Homology to the Human System"

```
                    10        20        30        40        50        60        70
HUMAN      MASAAAAEAEKGSPVVVGLLVVG-NIIILLSGLSLFAETIWVTADQYRVYPLMGVSGKDDVFAGAWIAIF 69
MOUSE      -MASAATEGEKGSPVVVGLLVVG-NIIILLSGLALFAETVWVTADQYRVYPLMGVSGKDDVFAGAWIAIF 68
ZEBRAFISH  ----------MGAVTCLMVTVVGLNAIAAAAGLALSAVAIWVAVDGYKLYPISGVSGKDDIFAGAWIAIF 60

                    80        90       100       110       120       130       140
HUMAN      CGFSFFMVASFGVGAALCRRRSMVLTYLVLMLIVYIFECASCITSYTHRDYMVSNPSLITKQMLTFYSAD 139
MOUSE      CGFSFFVVASFGVGAALCRRRYMILTYLLLMLIVYIFECASCITSYTHRDYMVSNPSLITKQMLTYYSAD 138
ZEBRAFISH  TGFAFFLTCIFGIFAALKRSRALMLVYLIIMFIIFLFESASAITSATNRDYLVGNSNLVKKQMLQYYADS 130

                   150       160       170       180       190       200       210
HUMAN      TDQGQELTRLWDRVMIEQECCGTSGPMDWVNFTSAFRAATPEVVFPWPPLCCRR-TGNFIPLNEEGCRLG 208
MOUSE      TDQGQELTRLWDRIMIEQECCGTSGPMDWVNYTSAFRAATPEVVFPWPPLCCRR-TGNFIPINEDGCRVG 207
ZEBRAFISH  STQGQQITMTWNNVMTQVQCCGADGPTDWIQYNSTYRQLFGAASL-WPLGCCKRQSSNFEVVDPIGCKAG 199

                   220       230       240       250       260
HUMAN      HMDYLFTKGCFEHIGHAIDSYTWGISWFGFAILMWTLPVMLIAMYFYTML- 258
MOUSE      HMDYLFTKGCFEHIGHAIDSYTWGISWFGFAILMWTLPVMLIAMYFYTTL- 257
ZEBRAFISH  VTSSMFTQGCFQYIESVLSRYTWAVSWYGFSVLMLVFFTLVIAMIYYTQLP 250
```

**Supplementary Figure 1. Alignment of Human, Mouse and Zebrafish Uroplakin 1a Amino Acid Sequences.** Muscle Tool used for alignment. Human sequence used as reference. Alignment highlighted in green.

```
                   10        20        30        40        50        60        70
HUMAN      MAPLLPIRTLPLILILLALLSPGAADFNISSLSGLLSPALTESLLVALPPCHLTGGNATLMVRRANDSKV  70
MOUSE      MASTLPVQTLPLILILLAVLAPGTADFNISSLSGLLSPALTESLLIALPPCHLTGGNATLMVRRANDSKV  70
ZEBRAFISH  -----------------------------------------------MPDCSIYGNQSVLLLYTEAPTNL  23

                   80        90       100       110       120       130       140
HUMAN      VTS-SFVVPPCRGRRELVSVVDSGAGFTVTRLSAYQVTNLVPGTKFYISYLVKKGTATESSREIPMSTLP  139
MOUSE      VKS-DFVVPPCRGRRELVSVVDSGSGYTVTRLSAYQVTNLTPGTKYYISYRVQKGTSTESSPETPMSTLP  139
ZEBRAFISH  NNTVNFTVQPCP-----------------VSQSWYLLGNLKNGTTYSMSYKI----GNDTSSVLTNTTTN  72

                  150       160       170       180
HUMAN      RRNMESIGLGM-ARTGGMVVITVLLSVAMFLLVLGFIIALALGSRK  184
MOUSE      RKNMESIGLGM-ARTGGMVVITVLLSVAMFLLVVGLIVALHWDARK  184
ZEBRAFISH  VNDYQQIDTGLRARSGAMVVITVILSLAMVFLLVGIILVFFFFSG-  117
```

**Supplementary Figure 2. Alignment of Human, Mouse and Zebrafish Uroplakin 2 Amino Acid Sequences.** Muscle Tool used for alignment. Human sequence used as reference. Alignment highlighted in green.

```
HUMAN      -----------MPPLWALLALGCLRFGSAVNLQPQLASVTFATNNPTLTTVALEKPLCMFDSKEA-----  54
MOUSE      -----------MLLLWALLALGCLRCGWTVNLQPQLASVTFATNNPTLTTVALEKPLCMFDSSEP-----  54
ZEBRAFISH  MNTHNAIRLVSLLSIWMLAAQGQI-------FQPQLAPANF-LGRITSNTVILQQPYCVFTQTCPGCEIW  62

HUMAN      -----LTGTHEVYLYVLVDSAISRNASVQDSTNTPLGSTFLQTEGGRTGPYKAVAFDLIPCSDLPSLDAI  119
MOUSE      -----LSGSYEVYLYAMVDSAMSRNVSVQDSAGVPLSTTFRQTQGGRSGPYKAAAFDLTPCGDLPSLDAV  119
ZEBRAFISH  LVAALSTGTGNFNALVNISSPISLSVSPYPTAFLPSSAQFFLT---RVGPLANF-----PCNTAPAF---  121

HUMAN      GDVSKASQILNAYLVRVGANGTCLWDPNFQGL-CNAPLSAATEYRFKYVLVNMSTGLVEDQTLWSDPIRT  188
MOUSE      GDVTQASEILNAYLVRVGNNGTCFWDPNFQGL-CNPPLTAATEYRFKYVLVNMSTGLVQDQTLWSDPIWT  188
ZEBRAFISH  -----------PYFT-VGADGIC------TGINCNGVLPVGSIVSFRYLLIDPSNYTVVNMTNWGGPFNL  173

HUMAN      NQLTPYSTIDTWPGRRSGGMIVITSILG-SLPFFLLVGFAGAIALSLVDMGSSDGETT----------HD  247
MOUSE      NRPIPYSAIDTWPGRRSGGMIVITSILG-SLPFFLLVGFAGAIILSFVDMGSSDGEMT----------HD  247
ZEBRAFISH  TTLLSYQTINDGLSARSGAMVVITTLLCVAVALLLLVFF---IMLCVSCCGKKDGKTVTVMSSIRIPRYD  240

HUMAN      SQITQEAV-PKSLGASESSYTSVNRGPPLDRAEVYSSKLQD  287
MOUSE      SQITQEAV-PKTLGTSEPSYSSVNRGPPLDRAEVFSSKLQD  287
ZEBRAFISH  THNLKEHVHPYDNQAYEPDAKNYSRSQTLPKSPVRK-----  276
```

**Supplementary Figure 3. Alignment of Human, Mouse and Zebrafish Uroplakin 3a Amino Acid Sequences.** Muscle Tool used for alignment. Human sequence used as reference. Alignment highlighted in green.

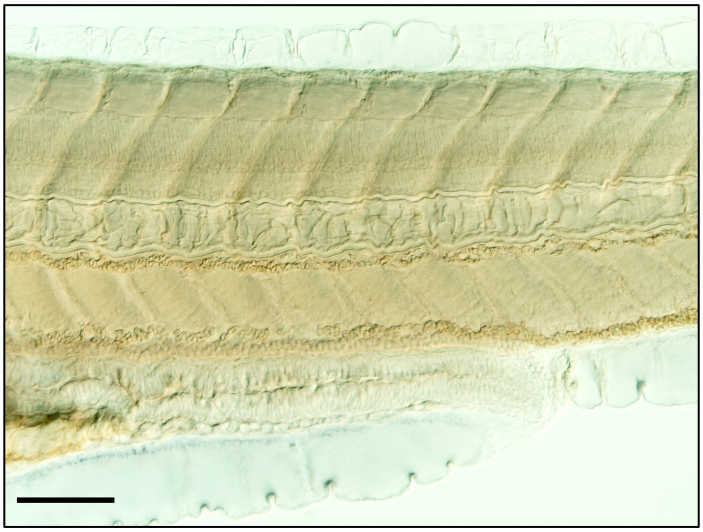

**Supplementary Figure 4. Sense Control for *Uroplakin 1a In Situ* Hybridisation in Zebrafish larvae at 96 hpf.** Scale bar= 50μm.

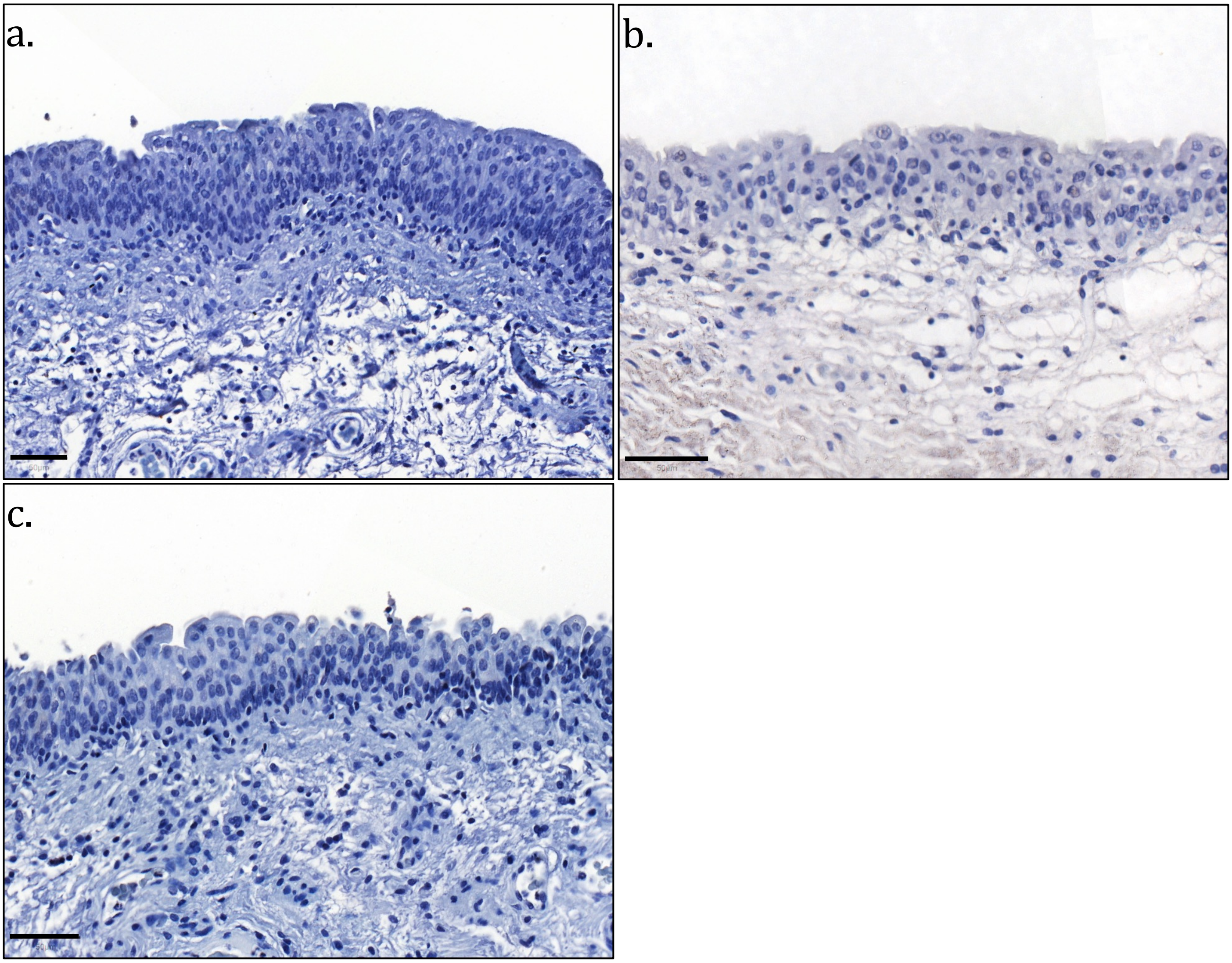


**Supplementary Figure 5. Negative (No Primary Antibody) Controls for Immunohistochemistry. A.** No primary antibody control for biotinylated goat anti-rabbit secondary antibody, **B.** No primary antibody control for biotinylated rabbit anti-goat secondary antibody. **C.** No primary antibody control for biotinylated goat anti-mouse secondary antibody. Scale bars= 50µm.
